# RodZ acts through MreBCD to activate the elongasome in *Escherichia coli*

**DOI:** 10.64898/2026.01.05.697639

**Authors:** Rui Zhan, Han Gong, Ying Li, Yuanyuan Cui, Xiangdong Chen, Joe Lutkenhaus, Shishen Du

## Abstract

Most bacteria are surrounded by a peptidoglycan (PG) matrix that maintains cell shape and provides protection against turgor pressure. In many rod-shaped bacteria, synthesis of PG along the cell cylinder is organized by the elongasome, also called the Rod complex, which consists of six highly conserved proteins, including the actin-like MreB, the PG synthase complex RodA-PBP2 and three regulatory membrane proteins, MreC, MreD and RodZ. However, how these proteins interact with each other to form the elongasome and synthesize lateral PG remains elusive. In this study, by characterizing MreC mutations affecting elongasome activity in *Escherichia coli*, we provide evidence that MreC alternates between an active and an inactive state and these mutations affect MreC interaction with PBP2. We also find that RodZ interacts with MreC via a periplasmic region and disruption of this interaction compromises elongasome function. Additionally, we show that the cytoplasmic region of RodZ, which interacts with MreB, work synergistically with activating MreC mutations to promote rod shape formation. These results indicate that RodZ activates the elongasome by initiating two signaling cascades, one in the periplasm via MreCD, and another in the cytoplasm through MreB. This role of RodZ in regulating elongasome activity is analogous to that of FtsN in activating the divisome, suggesting that the mechanisms governing the activation of PG synthesis in cell elongation and division are similar in *E. coli*.

**Importance:** The elongasome, or Rod complex, mediates lateral peptidoglycan synthesis during cell elongation in many rod-shaped bacteria. It consists of the cytoskeletal protein MreB, the peptidoglycan synthase RodA-PBP2, and three regulatory proteins MreCD and RodZ. Although it has been extensively studied, how its activity is controlled remains incompletely understood. Here, we reveal the roles of MreCD and RodZ in regulating elongasome activity in *Escherichia coli*. Our results indicate that RodZ triggers lateral peptidoglycan synthesis by interacting with MreB in the cytoplasm and MreCD in the periplasm. These interactions likely switch MreBCD to the active state such that they can stimulate the activity of the RodA-PBP2 complex. This regulatory mechanism resembles the activation mechanism of the divisome that mediates septal peptidoglycan synthesis.

## Introduction

The peptidoglycan (PG) layer determines the shape of bacterial cells and provides protection from osmotic lysis (1, 2). It is composed of glycan chains of repeated units of N-acetylglucosamine (GlcNAc) and N-acetylmuramic acid (MurNAc), which are interconnected by short peptides bridges, resulting in a continuous matrix enveloping the bacterial membrane (1–3). Biosynthesis of the PG layer starts with the synthesis of its precursor lipid II, a disaccharide pentapeptide embedded in the cytoplasmic membrane by attachment to the lipid carrier undecaprenyl phosphate (Und-P). Lipid II is then transported to the outer face of the cytoplasmic membrane, where it is polymerized into glycan chains by PG glycosyltransferases (GTase) and then crosslinked to the existing PG network by transpeptidases (TPase) (1, 2). Class A penicillin-binding proteins (aPBPs), which harbor both GTase and TPase activities, were previously believed to be the main PG synthases in bacteria (1, 2). However, recent studies demonstrated that cognate pairs of SEDS (shape, elongation, division, spore formation) proteins and class B PBPs (bPBPs) constitute the main PG synthases for PG expansion and septation, whereas the aPBPs fortify weak areas of the PG layer to maintain its integrity (4–10). Inhibition of PG biosynthesis results in cell shape and division defects, ultimately leading to cell lysis. An in-depth understanding of the mechanism of PG biosynthesis is expected to aid the development of new inhibitors of cell wall synthesis (11).

Most rod-shaped bacteria employ two multi-protein complexes, the elongasome and the divisome, to assemble the PG matrix (2). The elongasome, or Rod complex, is responsible for the synthesis of lateral PG to promote cell elongation, while the divisome is dedicated for the synthesis of septal PG (sPG) during cell division. In *Escherichia coli,* the elongasome consists of six highly conserved proteins, MreB, MreC, MreD, RodA, PBP2 and RodZ, whereas the divisome contains 10 essential and many accessory components, including the tubulin-like FtsZ and actin-like FtsA (1, 2, 12). Despite the difference in composition, the two complexes are evolutionary related as some of their components belong to the same protein families (13). For example, both MreB and FtsA are actin-like proteins, and both complexes contain a SEDS-bPBP PG synthase complex (RodA-PBP2 *vs* FtsW-FtsI) (10, 14). However, the proteins involved in the regulation of the SEDS-bPBP complexes display no homology (MreCD-RodZ for the elongasome and FtsQLBN for the divisome) raising questions about the regulatory mechanisms. Understanding how these protein complexes orchestrate PG synthesis during cell elongation and division has been and continues to be one of the research focuses of PG biology.

The elongasome has been extensively studied, but its regulatory mechanism is not fully clear. MreB assembles into anti-parallel double filaments anchored to the inner leaflet of the cytoplasmic membrane, which organize the assembly and activity of the elongasome complex (15–20). Consequently, disruption of MreB assembly by certain MreB mutations or by the small molecule A22, which is a potent MreB inhibitor, results in a block of lateral PG synthesis and a loss of rod shape (21–23). Intriguingly, MreB filaments display a directional movement perpendicular to the long axis of the cell, which is driven by ongoing lateral PG synthesis rather than its ATPase activity (19, 20, 24). It has been suggested that MreB filaments function in a rudder-like mechanism to guide lateral PG synthesis (16), but the exact mechanism is still enigmatic. RodZ is a bitopic membrane protein with a cytoplasmic helix-turn-helix (HTH) domain, which binds MreB, and a periplasmic domain, which interacts with the other components of the elongasome (25–29). The deletion of *rodZ* causes the formation of aberrant MreB structures and mislocalization of the other elongasome components, ultimately leading to a partial loss of rod shape and conditional growth (25, 30). Thus, RodZ likely modulates the assembly of MreB filaments and couples them with other components of the elongasome (30, 31). RodA is a member of the SEDS family of PG polymerases (5, 6), which partners with PBP2 (a member of bPBPs) to carry out PG synthesis by the elongasome (7, 10). The structure of RodA-PBP2 has been resolved and the PG polymerase activity of RodA relies on the presence of PBP2 (9, 10, 32), but a detailed regulatory mechanism of their enzymatic activity awaits elucidation. MreC and MreD form a tight complex with MreB and recent studies indicate that they play an essential role in regulating RodA-PBP2 activity (9, 33, 34).

A picture emerging from the studies of the elongasome and divisome complexes over the last decade is that the activity of the PG synthases (RodA-PBP2 and FtsW-FtsI) within them are tightly regulated to ensure the expansion and division of the PG matrix occur in the right place at the right time (2, 9, 14)(Fig. 1). During division, the essential protein FtsN triggers sPG synthesis by initiating two signaling cascades, one in the cytoplasm and one in the periplasm (35–37). The cytoplasmic and periplasmic domains of FtsN switch FtsA and the FtsQLB subcomplex to an active state, respectively, such that the active conformer of the FtsWI complex is stabilized, leading to sPG synthesis (14, 35–40). Interestingly, mutations in the extracellular loop 4 (ECL4) of FtsW and in the pedestal domain of FtsI, which constitutes their second interaction interface, bypass the requirement of FtsN for sPG synthesis (14, 41). Strikingly, mutations in similar regions of RodA and PBP2 result in constitutively active RodA-PBP2 complex, indicating an essential role for the interaction interfaces in modulating the activity of these SEDS-bPBP complexes (9, 42). Structural and biochemical analysis of the RodA-PBP2 complex suggest that it transitions between closed and open states, which likely represent the inactive and active state of the complex, respectively (10, 43). A structure of the *Helicobacter pylori* MreC-PBP2 complex shows that the periplasmic domain of MreC interacts with the pedestal domain of PBP2, resulting in conformational changes in PBP2 (44), suggesting that MreC binding switches the RodA-PBP2 complex to the active state. Consistent with this, certain mutations in MreC (G156D or R292H) result in the inactivation of the elongasome, but the above mentioned constitutively active mutations in PBP2 and RodA suppress these MreC mutations and restore rod shape (9). MreC forms dimers or double antiparallel filaments, which would occlude its PBP2 binding domain (45–47). Thus, it is believed that modulation of MreC self-interaction is critical for its interaction with PBP2 and stabilization of the active conformation of the RodA-PBP2 complex (10, 46, 48). However, how MreC transitions between the two states is unclear and direct evidence for MreC(D) controlling RodA-PBP2 activity is still lacking.

**Fig. 1.**
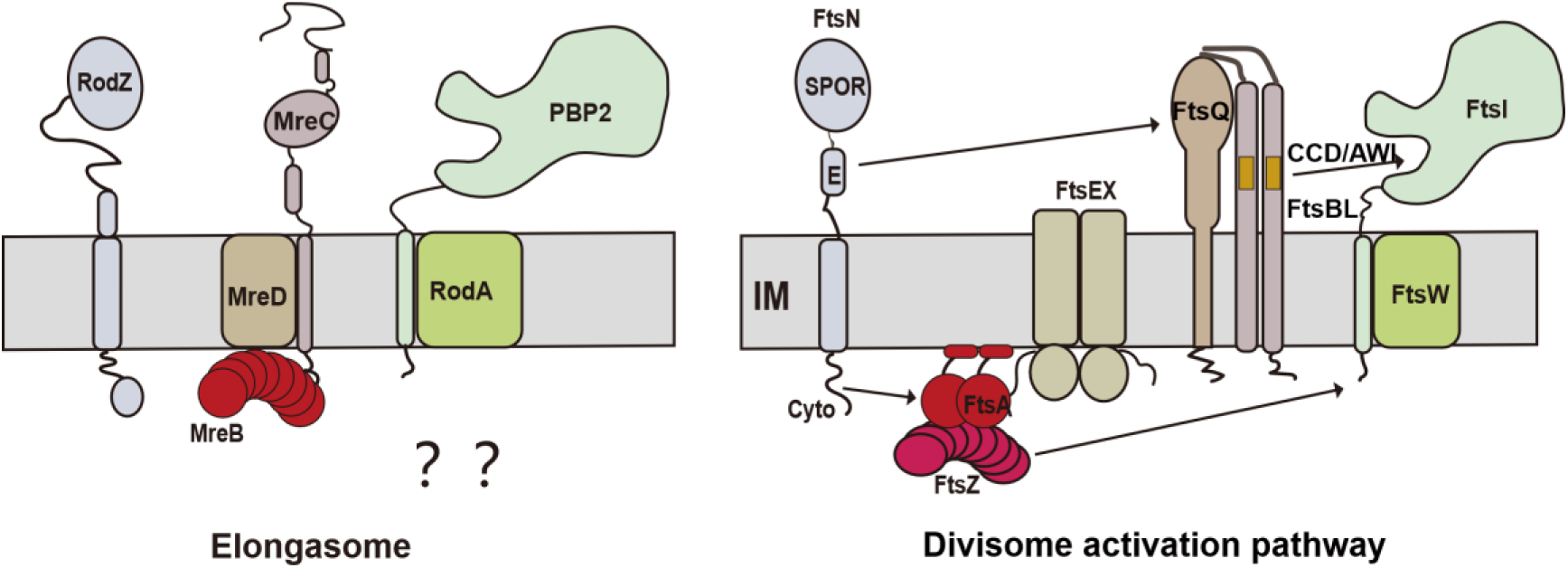
Activation pathways of the elongasome and divisome. Left: The elongasome is composed of six proteins: MreB, MreC, MreD, RodA, PBP2 and RodZ. However, the regulatory mechanism for the activation of elongasome is not yet fully understood. Right: During division, the essential protein FtsN triggers sPG synthesis by initiating two signaling cascades, one in the cytoplasm and the other in the periplasm. The cytoplasmic and periplasmic domains of FtsN switch FtsA and the FtsQLB subcomplex to an active state, respectively, such that the active conformer of the FtsWI complex is stabilized, leading to sPG synthesis. The CCD/AWI domains of FtsB and FtsL play important roles in regulating FtsW-FtsI activity. CCD: constriction control domain; AWI: activation of FtsWI domain.

In this study, we isolated MreC mutations that increase the activity of the elongasome in *E. coli* and found that many of them enhance the interaction between MreC and PBP2, but there is no clear correlation with its oligmerization state. We identified an interaction between RodZ and MreC in the periplasm, which appears to be important for elongasome function. Moreover, the cytoplasmic domain of RodZ and activating MreC mutations work synergistically to promote rod-shape formation. Taken together, our results suggest a model in which RodZ functions as a trigger for the elongasome. In the periplasm, RodZ switches MreC to the active state so that it binds PBP2 and stabilizes RodA-PBP2 in an active conformation; in the cytoplasm, RodZ interacts with MreB, leading to the activation of RodA-PBP2 via a yet unknown mechanism. This activation pathway resembles the activation mechanism of the divisome, suggesting a shared regulatory framework in governing PG synthesis during cell elongation and division.

## Results

### Isolation of *E. coli* MreC mutants that enhance elongasome activity

Characterization of dominant-negative (also inactive) MreCD mutations suggested that MreCD alternates between an inactive and an active state (34). We hypothesized that certain mutations in MreCD may stabilize the complex in the active state and enhance elongasome activity. In-depth analysis of such mutations may provide important insights into its working mechanism. Because previous studies have found that increasing the integrity and/or the activity of the elongasome confers resistance to A22, an MreB antagonist (22, 42, 49), we screened for MreCD mutants that rescued cell growth in the presence of a lethal concentration of A22. To do this, we constructed a plasmid library encoding a PCR-mutagenized version of the *mreCD* operon under the control of an IPTG (isopropyl-β-D-thiogalactopyranoside)-inducible promoter. The library was then transformed into a wild-type strain (JS238) and transformants were selected on LB plates containing 30 µM IPTG and 10 μg/mL of A22. A22-resistant transformants arose at a relatively high frequency (10^-4^) and 16 were randomly picked for further study. Plasmids in these transformants were isolated and retransformed into the wild type strain to confirm that the A22-resistance was due to the plasmids (Supplementary Fig. 1a). Sequencing of *mreCD* in these plasmids revealed that they harbored one or several mutations (Supplementary Table 1). To determine the causative mutations for resistance to A22, single substitutions were introduced into the parental plasmid and assayed for A22 resistance. Strikingly, we found that 11 *mreC* mutations (E100D, P150L, S153R, D154N, P182L, I183F, Q184L, T222A, E231K/V and R264H), but none of the *mreD* mutations, provided A22-resistance (Fig. 2a and Supplementary Fig. 1b), suggesting that MreC plays a prominent role in regulating elongasome activity. Interestingly, one plasmid harboring a single mutation in *mreC* (G259V) and one in *mreD* (L50F) conferred A22-reistance, however, neither mutation alone could confer resistance (Supplementary Fig. 1b), suggesting a coordination between MreC and MreD in regulating elongasome activity.

**Fig. 2.**
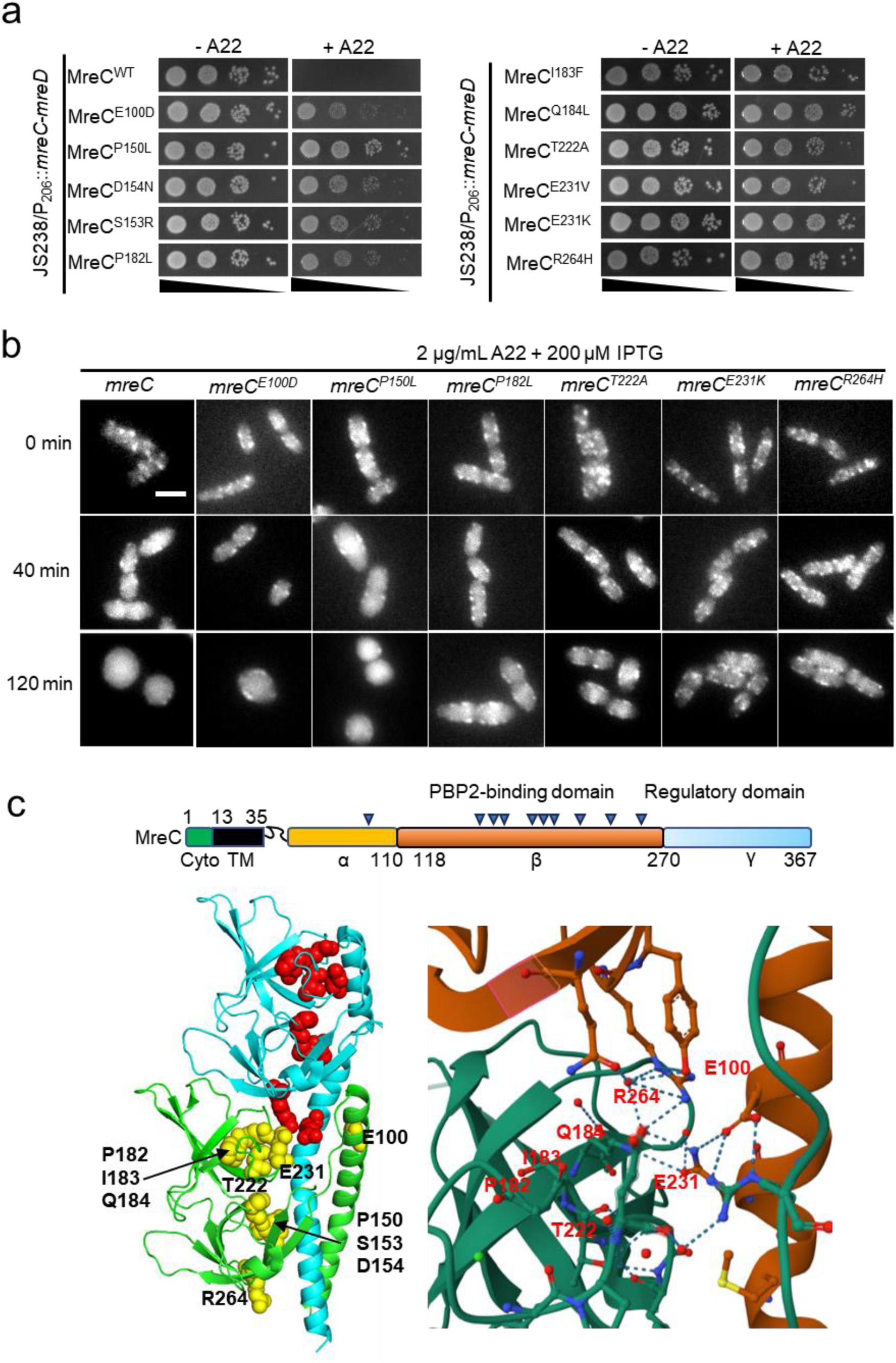
Isolation of MreC mutants that enhance elongasome activity in *E. coli*. (a) Mutations in *mreC* provide resistance to the MreB antagonist A22. Plasmids pSD315 (P_206_::*mreC-mreD*) harboring the indicated *mreC* mutations were transformed into strain JS238. Resistance of the transformants to A22 was assessed on LB plates with antibiotics, with or without A22 (10 µg/mL) and 30 µM IPTG. Plates were incubated at 37°C overnight and photographed. (b) Representative images of MreB localization (^SW^MreB-mNeonGreen: *^SW^mreB-mNG*) and cell shape of the indicated strains with or without A22 treatment. Strains (RZ15 to RZ21) expressing *^SW^mreB-mNG* were grown to log phase and 2 μg/mL A22 was added to the culture (time 0). Samples were harvested at 0 min, 40 min or 120 min after the addition of A22 and observed. Induction of *^SW^mreB-mNG* (from attλHC897) was achieved with 200 μM IPTG. (c) Locations of the mutations in the structure of *E. coli* MreC dimer (PDB ID#:7EFT). A diagram showing the five subdomains of *E. coli* MreC is shown above the structure. Cyto (cytoplasmic region, 1-13), TM (transmembrane segment, 14-35), α (36-110), β (111-270) and γ (271-367) domains. Amino acid substitutions conferring resistance to A22 are indicated by triangles in the diagram and highlighted in yellow or red in the MreC dimer structure (PDB ID#:7EFT).

To further characterize the *mreC* mutations, 9 of them were successfully introduced into the chromosome by allelic replacement (50). 5 of the resultant strains displayed modest to strong A22 resistance (S153R, D154N, T222A, E231K and R264H), but the rest exhibited low resistance (E100D, P182L and Q184L) or even became more sensitive (P150L) compared to wild type strain (Supplementary Fig. 1c), implying that the second set of mutants may need to be overexpressed or rely on the presence of wild type MreC to confer A22 resistance. Several of these mutants became more sensitive to the PBP2 inhibitor mecillinam (Supplementary Fig. 1d), similar to the previously reported PBP2^L61R^ (33). MreB localization and cell morphology in a subset of these strains were also examined after A22 treatment. As shown in Fig. 2b, MreB localization in wild type cells switched from the typical spotty membrane foci into evenly distributed cytoplasmic fluorescence by 40 min post A22 treatment and cells became completely round by 2 hours. However, in cells harboring *mreC* mutations providing strong resistance to A22 (T222A, E231K and R264H), rod-shape was still partially maintained at 2 hours after A22 treatment and MreB foci could still be observed at this time. The phenotypes of these *mreC* mutations are similar to activating RodA and PBP2 mutations (42), implying that they enhance the integrity/activity of the elonagsome. As expected, cells carrying *mreC* mutations exhibiting low resistance or no resistance (E100D, P150L and P182L) behaved more or less like wild type cells in terms of MreB localization and morphology (Fig 2b).

Since previous studies found that mutations enhancing elongasome activity alleviate the growth and shape defects of *ΔrodZ* cells (9, 42, 49), we tested whether the isolated *mreC* mutations had a similar effect. 6 out of the 9 tested *mreC* mutations, including E100D, P182L, Q184L, T222A, E231K and R264H, ameliorated the viability defects of *ΔrodZ* cells upon shifting the strains from 30°C (permissive condition) to 22°C (non-permissive condition), accompanied by partial restoration of rod shape (Supplementary Fig. 2a-b). However, three *mreC* mutations (P150L, S153R and D154N), failed to rectify the growth and shape defect of *ΔrodZ* cells (Supplementary Fig. 2a-b), suggesting that they may depend on RodZ to exert their effect.

Lastly, we tested if the isolated MreC mutations could function as intragenic suppressor of the previously reported dominant-negative MreC mutation (G156D and R292H) (9, 34). These two mutations were believed to lock MreC in the inactive state, thus resulting in elongasome inactivation. Intriguingly, most of the tested mutations function as intragenic suppressor of MreC^R292H^ (except for the E100D), suggesting that they relieve the inhibitory effect of the R292H mutation. However, none of the mutations suppressed MreC^G156D^ (Supplementary Fig. 2c-d), indicating that G156D may inactivate MreC by a different mechanism than that of R292H.

Based on the results of A22 sensitivity test and the ability to suppress the growth and shape defects caused by *rodZ* deletion or *mreC* inactive mutations, we conclude that most of our *mreC* mutations enhance elongasome activity, but their strength and underlying mechanisms might be different. Also, some appear to rely on the presence of RodZ to display their effect, while others do not. We termed mutations increasing integrity/activity of the elongasome activating mutations, while mutations reducing or blocking elongasome activity are designated inactivating mutations in the rest of the paper.

### Activating MreC mutations cluster at two distinct regions

*E. coli* MreC can be artificially divided into five subdomains, a short cytoplasmic region (1–13), a transmembrane segment (14–35), followed by a helical α domain (36–110), a β domain consisting of mainly β strands (111–270) and a proline-rich γ domain (271–367) (Fig. 2c). Most of the mutated residues are in the β domain of MreC, except for E100, which is in the α domain (Fig. 2c). Interestingly, most of these residues are conserved in Gram-negative bacteria, but not in Gram-positive bacteria (Supplementary Fig. 3), implying a different regulatory mechanism of MreC function in these two kinds of bacteria. Mapping the mutated residues onto the dimer structure of the periplasmic domain of *E. coli* MreC (PDB#: 7EFT) revealed they mostly clustered in two regions. The first set of mutations, including E100, P182L, I183F, Q184L, T222A, E231K and R264H, are at the homo-dimer interface between the α and β domains of the two monomers (Fig. 2c, yellow). The other set of mutations (P150L, S153D and D154N) are located in one of the β-barrels of the β domain (Fig. 2c), whose corresponding residues are not at the MreC dimer interface. However, mutations in neighboring residues (K155E and G156D) inactivate MreC (51), suggesting that this region plays an important role in fine tuning MreC activity. It is notable that the second set of mutations (P150L, S153R, and D154N) appeared to rely on RodZ to exert their effects (Fig. 2c), implying that there is a genetic interaction between MreC and RodZ in regulating elongasome activity. Taken together, the above analysis suggests that there are two regions in MreC that are critical for it to control elongasome activity in addition to its interaction site with PBP2 and the γ domain, one in the self-interaction interface, the other within an internal region of the β domain.

### There is no clear correlation between MreC self-interaction and its functional states

To investigate how the *mreC* mutations affect its function, we employed a bacterial-two-hybrid (BTH) assay to test whether they affect MreC self-interaction, because recent studies suggested that self-association of MreC precludes it from interacting with PBP2 (46). As shown in Supplementary Fig. 4, MreC mutants behaved similarly as wild type MreC, displaying strong self-interaction and interaction with PBP2, indicating that the mutations either do not affect MreC self-interaction or the assay could not distinguish the mutants from wild type MreC because of the presence of other elongasome components.

To further investigate how the activating mutations influence MreC activity, we employed BMOE-mediated *in vivo* cysteine-crosslinking assay, which has been widely used to access protein-protein interactions and protein conformational changes *in vivo*. In this assay, the thio-reactive reagent BMOE crosslinks two cysteine residues within a distance of ∼8 Å (15), resulting in the formation of cross-linked species (CLS). We introduced two cysteine substitutions, L108C and Q122C, in the dimerization interface of MreC (Fig. 3a). The Cα-Cα distances between the two L108C residues in the MreC dimer structure was 6.7 Å, within the working distance of BMOE, whereas the Cα-Cα distances between the Q122C residues or between L108C and Q122C were longer than 8 Å. We expected that the L108C mutant would generate a dimeric product in the presence of BMOE while the Q122C mutant would not. Complementation test showed that neither cysteine mutation affected the ability of MreC (in tandem with MreD) to complement the depletion strain (Supplementary Fig. 5a). In the presence of BMOE, we detected a band with a molecular weight corresponding to an MreC dimer with the L108C mutant (Fig. 3a), however, we did not detect any crosslinked product for the Q122C mutant. This suggested that MreC^L108C^ but not MreC^Q122C^ could be utilized for testing the effect of mutations on MreC self-interaction or conformational changes. We first tested if the activating mutations retained the ability to complement and confer A22 resistance in the L108C mutant background. Many of the mutants retained the ability to complement the depletion strain, but only E231K and R264H in the background of L108C could provide weak resistance to A22 (Supplementary Fig. 5b-c). Thus, we examined how these two mutations influenced the crosslinking of MreC^L108C^ in the presence of BMOE. Interestingly, E231K drastically reduced the crosslinked product, whereas R264H had no effect (Fig. 3b-c), indicating that E231K disrupts MreC self-interaction or causes a conformational change in the dimerization interface of MreC, but the R264H mutation might shift MreC to the active state by affecting other aspects of MreC.

**Fig. 3.**
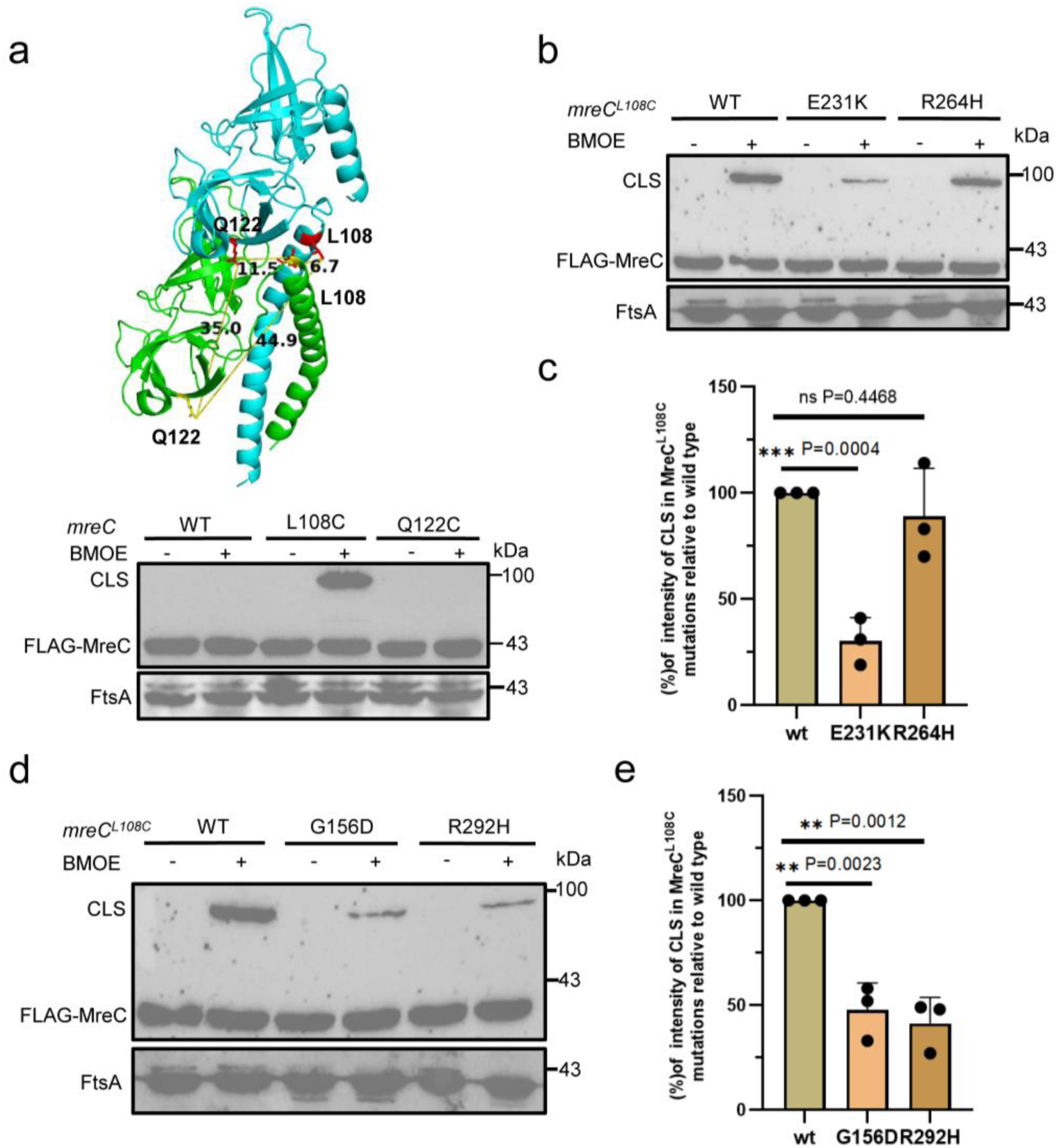
Impact of activating and inactivating MreC mutations on its self-interaction and conformation. (a) Locations of the introduced cysteine residues on the MreC dimer structure (PDB#: 7EFT). Cysteine L108C and Q122C were chosen to test MreC self-interaction or conformational change by BMOE-mediated *in vivo* cross-linking. The Cα-Cα distance between the cysteine pairs was marked in the legend. Samples were obtained from *mreCD* knockout strain SD464 (TB28, *mreCD<>aph*, P_BAD_::*mreC-mreD*) supplemented with a cysteine-containing *flag-mreC* allele (P_206_::*flag-mreC^C175S,C199S^-mreD*), with mutations labeled at the top of each lane. Cells were grown until the OD_600_ reached 0.4 to 0.6, an aliquot of the culture was collected and washed twice in PBST. The suspension was split into two halves, one treated with (+) BMOE, and the other half without (-). Cells were then incubated on ice for 15 minutes, followed by the addition of β-BME to terminate the cross-linking reaction. Samples were subsequently analyzed by Western blotting using a Flag-tag antibody. FtsA was used as a loading control. (b-c) Different effects of activating mutations on MreC self-interaction as determined by BMOE-mediated cross-linking assay. The assay was carried out as in (a). Representative plot for the results from three independent experiments was shown in (b) and comparison of crosslinking efficiency (mean±standard error of the mean (SEM)) based on the intensity of crosslinked products relative to wild type MreC was shown in (c). (d-e) Inactivating mutations disrupt MreC self-interaction as determined by BMOE-mediated crosslinking assay. The experiment was performed as in (b). Representative plot for the results from three independent experiments was shown in (f) and relative crosslinking efficiency against wild type MreC was shown in (g). Statistical significance was determined using an unpaired *t* test with Welch’s correction in (d) and (g), ns: non-significant.

Since the effect of activating mutations on MreC self-interaction or conformation was promiscuous, we wondered how inactivating MreC mutations influenced the its self-interaction or conformation, such as the two well characterized dominant-negative mutations, MreC^G156D^ and MreC^R292H^. BMOE-mediated *in vivo* crosslinking assay showed that the crosslinked MreC^L108C^ dimer was markedly reduced in the presence of the G156D or R292H mutation (Fig. 3d-e). It appeared that inactivation of MreC might involve disruption of the MreC self-interaction or conformational changes in the interfaces, but activation of MreC does not necessarily require an alteration of MreC self-interaction. Taken together, we conclude that there is no clear correlation between the oligomerization state of MreC and its activity in *E. coli*.

### Activating and inactivating MreC mutations affect the interaction between MreC and PBP2

Since the interaction between MreC and PBP2 is critical for regulating elongasome activity, we wondered if the activating MreC mutations enhanced MreC interaction with PBP2. MreC binds to PBP2 mainly through its β domain (44), but the γ domain of MreC (aa279-367) is believed to regulate this interaction as mutations, such as R292H, in a short helix (279–305) within this domain, inhibits elongasome activity (34). However, the defects of MreC^R292H^ can be suppressed by activating mutations in PBP2/RodA and a truncated form of MreC (MreC^1-278^) complemented the MreC depletion strain. It had been suggested that the γ domain regulated MreC activity by an auto-inhibitory mechanism (34). We found that MreC variants lacking parts of or the entire γ domain, MreC^1-305^ or MreC^1-278^ (in tandem with MreD), failed to complement the MreCD depletion strain at low levels of induction, but did at higher levels of induction (Supplementary Fig. 6a). Western blotting of the protein level of the truncated forms of MreC (1-305 and 1-278) fused to GFP showed that their levels were comparable but lower than that of full-length MreC (Supplementary Fig. 6b). However, MreC^1-305^ appeared to be slightly stronger than MreC^1-278^ in complementation (Supplementary Fig. 6a), suggesting the region (aa279-305) may contribute to MreC activity. In line with this, we found that MreC^1-305^ displayed a decreased interaction with PBP2 in comparison to wild type MreC in the Bacterial Two Hybrid (BTH) assay, while MreC^1-278^ failed to interact with PBP2 (Supplementary Fig. 6c). Nonetheless, both mutants retained the ability to interact with other elongasome components. Thus, we decided to compare the binding strength of MreC^1-305^ and MreC^1-278^ with PBP2. Co-immunoprecipitation experiments showed that GFP-tagged MreC^1-278^ interacted with PBP2 less well than GFP-tagged MreC^1-305^ (Supplementary Fig. 6d). These results indicate that the γ domain is necessary for the optimal interaction between MreC and PBP2, especially the motif consisting of residues 279-305. In the absence the γ domain, MreC (MreC^1-278^) retains the ability to interact with PBP2 via it β domain but is weaker.

The above observations prompted us to examine whether the activating *mreC* mutations could restore the interaction between MreC^1-278^ and PBP2. Strikingly, introduction of any one of the activating mutations in MreC^1-278^ restored the interaction in the BTH assay (Fig. 4a). Consistently, Co-IP experiments showed that introduction of either E231K or R264H, two representative activating mutations, into MreC^1-278^ increased its interaction with PBP2 (Fig. 4b-c). Consequently, while MreCD depleted cells expressing MreC^1-278^ (with MreD) displayed a rod shape defect, the presence of the activating mutations greatly improved the rod shape morphology and growth viability (Fig. 4d and Supplementary Fig. 7a). Western blotting suggested that the difference was not due to variation in protein level (Supplementary Fig. 7b). Altogether, these results suggest that MreC^1-278^ is unable to interact with PBP2 efficiently, but activating mutations promote MreC’s interaction with PBP2, thereby suppressing its defect and leading to increased elongasome activity (Fig. 4e).

**Fig. 4.**
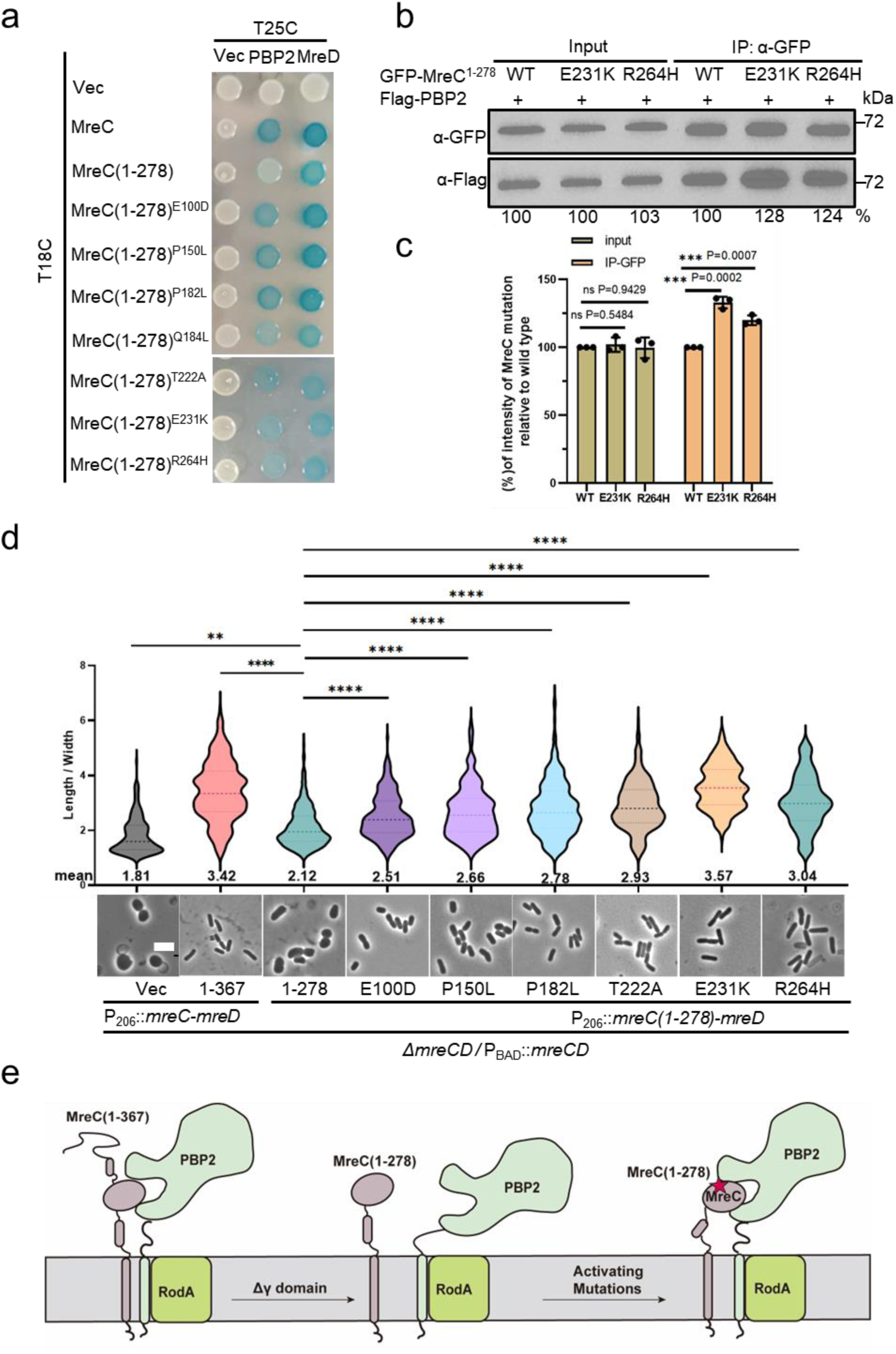
Activating MreC mutations enhance the interaction between MreC and PBP2. (a) Examination of the effect of activating MreC mutations on the interaction between MreC^1-278^ and PBP2 by BTH. Plasmid pairs were co-transformed into BTH101 and assay performed as in Fig. 3a. Interaction between T18C-MreC^1-278^/T25C-MreD served as a positive control. (b-c) Examination of the effect of activating MreC mutations on the interaction between GFP-tagged MreC^1-278^ and Flag-tagged PBP2 by Co-IP. Cells expressing the indicated forms of GFP-MreC^1-278^ and Flag-PBP2 in exponential phase were harvested and lysed by sonication. After removal of the debris, supernatants were collected and incubated with anti-Flag or Anti-GFP antibodies coated beads. Immunocomplexes were isolated and analyzed by western blotting. Intensities of the bands were quantitated by Image J. A representative plot for three independent experiments was shown in (b). Comparison of the amount of proteins in the immunocomplexes was based on the intensity of protein bands (c). Statistical significance was determined using an unpaired *t* test with Welch’s correction, ns: non-significant). (d) Activating MreC mutations suppress the shape defect caused by MreC^1-278^. Plasmids expressing the indicated MreC(D) mutants were transformed into an MreCD depletion strain. Overnight cultures of the strains expressing *mreC^1-278^(D)* or its variants were collected by centrifugation and washed twice with fresh LB to remove arabinose and resuspended in the same volume of LB supplemented with antibiotics and 60 µM IPTG and grown at 37°C. When the OD_600_ reached 0.4 to 0.6, cells were fixed and imaged. The aspect ratio (length/width) of cells were quantified using Image J. The ratio calculated for each cell is plotted as a violin dot, showing the 25th and 75th percentiles, with the mean at the center, whiskers extend from -s.d. to +s.d. of the mean. Number of cells analyzed (n>200). Statistical significance determined using an unpaired *t* test with Welch’s correction. Scale bar, 5 µm. **P < 0.01, ****P < 0.0001, two-tailed Student’s t-test. (e) Summary of the effects of the γ domain and activating MreC mutations on the interaction between MreC and PBP2. Left panel: full-length MreC is able to switch to the active form to interact with PBP2; middle panel: deletion of the γ domain (MreC^1-278^) prevents MreC from transitioning to the active form effectively, weakening the MreC-PBP2 interaction; right panel: activating *mreC* mutations convert MreC^1-278^ to the active form, restoring the MreC-PBP2 interaction.

Because the inactivating MreC mutations have an opposite influence on elongasome activity, we tested if they reduced the interaction between MreC and PBP2. As shown in Supplementary Fig. 8a, the inactive R292H mutation did not substantially reduce the interaction between MreC^1-305^ and PBP2 in the BTH assay, but the G156D mutation abolished the interaction signal. Moreover, Co-IP experiments showed that that both the G156D and R292H mutations reduced the interaction between MreC^1-305^ and PBP2 (Supplementary Fig. 8b-c). Thus, these mutations likely inactivate MreC by reducing its ability to interact with PBP2.

### An interaction between RodZ and MreC is necessary for elongasome activity

The results above indicate that MreC(D) switches from an inactive to an active state to interact with PBP2, but what triggers the transition has not been determined. A likely candidate is RodZ because it has been shown to interact with MreC in a BTH assay (25). Moreover, the shape defect of Δ*rodZ* cells is partially suppressed by certain MreC activating mutations, and several activating MreC mutations appear to depend on RodZ to exert their effect (Supplementary Fig. 2a-b). Thus, we hypothesized that an interaction between RodZ and MreC in the periplasm switches MreC to an active state (Fig. 5a). Previous studies suggested that the periplasmic domain of RodZ is dispensable for its function since truncated GFP-RodZ variants lacking most or all of the periplasmic domain (GFP-RodZ^1-155^ and GFP-RodZ^1-138^) largely restore growth and rod-shape morphology to *ΔrodZ* cells (25, 26). However, when we assessed the ability of these two GFP-RodZ variants on cell growth and shape using an expression vector with the arabinose promoter (P_ara_), we found that GFP-RodZ^1-155^, but not GFP-RodZ^1-138^, was able to complement the Δ*rodZ* strain at low levels of the inducer arabinose from 0 to 0.1% (Fig. 5b and Supplementary Fig. 9a). Western blotting showed that the protein levels of GFP-RodZ^1-155^ and GFP-RodZ^1-138^ were comparable (Supplementary Fig. 9b), indicating that the difference was due to the lacking of a periplasmic motif (aa 138–155) of RodZ. Nonetheless, both were able to rescue the growth defect of *ΔrodZ* strain at 0.2% arabinose (Fig. 5b). It is possible that the previous study (25) conducted the test by expressing GFP-RodZ^1-155^ and GFP-RodZ^1-138^ from an IPTG-inducible P_lac_ promoter, resulting in overexpression of these truncated forms of RodZ and masking the importance of the RodZ^138-155^ motif.

**Fig. 5.**
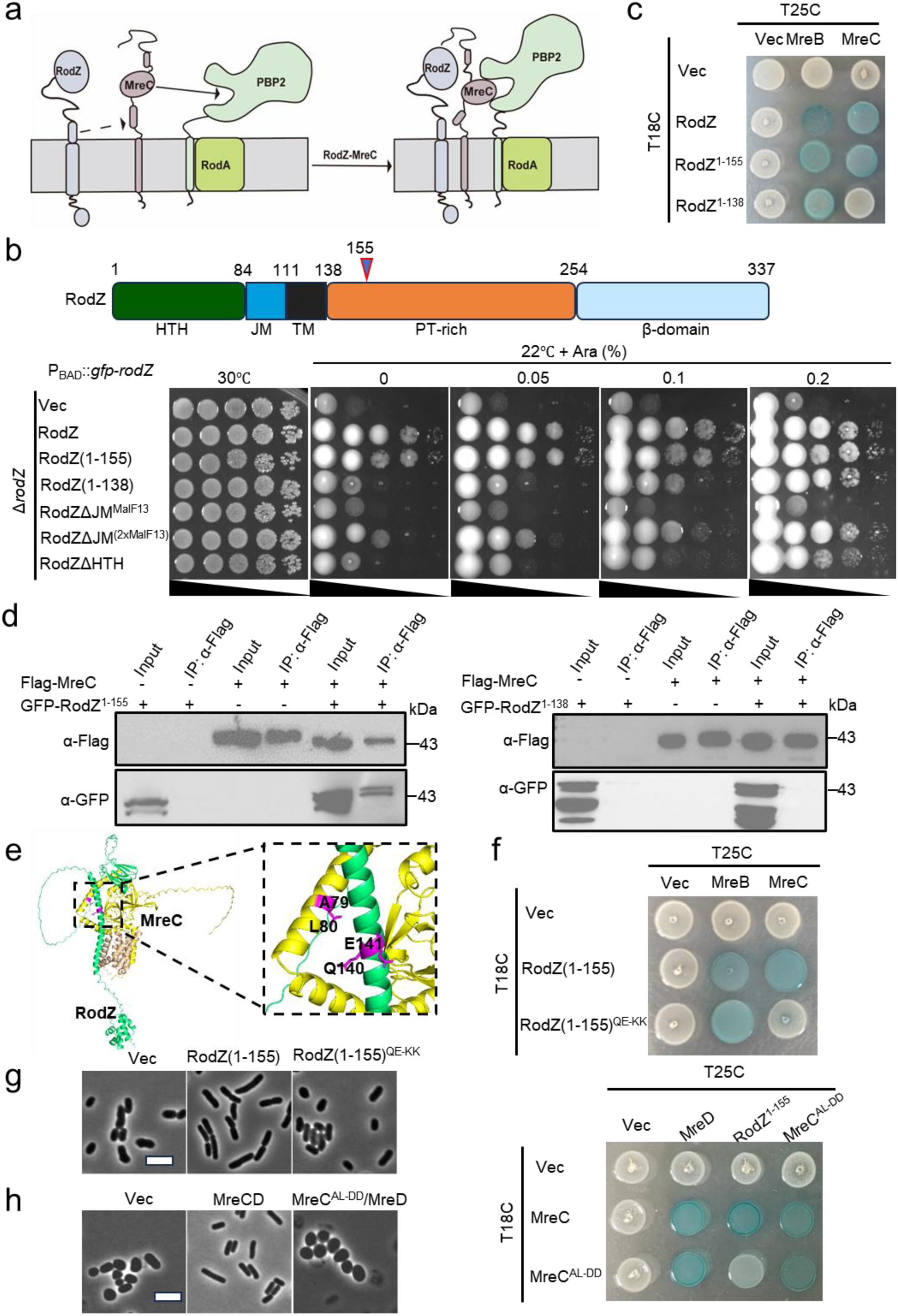
Interaction between RodZ and MreC is necessary for elongasome activity. (a) A hypothetical model for the activation of MreC by RodZ. A periplasmic domain of RodZ interacts with MreC, switching it from the inactive state to the active state such that it can interact with PBP2 to activate lateral peptidoglycan synthesis. (b) Schematic diagram of RodZ and characterization of its mutants. RodZ can be artificially divided into five domains as indicated. HTH: Helix-Turn-Helix; JM: juxta-membrane; TM: transmembrane; PT-rich: proline and threonine-rich region; β-domain: domain consisting of β-strands. Overexpression of RodZ^1-155^, but not RodZ^1-138^, rescued the growth of the Δ*rodZ* strain. Plasmid pBAD18, pZR102(P_BAD_::*gfp-rodZ*) or its derivatives were transformed into strain SD541 (TB28, *yhdE::cat*, *rodZ*<>*aph*) on LB plates at 30°C. Transformants were resuspended, serially diluted and spotted on plates with or without arabinose. Plates were incubated at 22°C for 40 hours or at 30°C overnight and photographed. The *rodZ* alleles include *gfp*-*rodZ^1-155^*, *gfp-rodZ^1-138^*, *gfp-rodZΔJM^(malF13)^*, *gfp-rodZ*Δ*JM^(^*^2x*malF13)*^ and *gfp-rodZ*Δ*HTH.* (c) RodZ^1-155^ but not RodZ^1-138^ interacts with MreC in BTH assay. Plasmid pairs were transformed into strain BTH101 and transformants were spotted on LB plates containing antibiotics, 40 μg/mL X-gal and 100 μM IPTG. Plates were incubated at 30°C for about 16 hours before photographing. (d) RodZ^138-155^ is critical for the interaction between RodZ and MreC. Overnight cultures of TB28 harboring plasmids pZR269 (pDSW209, *bla* P_206_::*gfp-rodZ^1-155^*-*flag-mreC*) and pZR270 (pDSW209, *bla* P_206_::*gfp-rodZ^1-138^*-*flag-mreC*) were diluted 1:100 in 50 mL fresh LB medium, grown at 37°C until OD_600_ =0.4-0.6. Samples were prepared for Co-IP experiments as described in Methods. (e) AlphaFold 3 model of ^Ec^RodZ-MreCD complex reveals residues critical for their interaction (RodZ^138-155^; MreC^78-83^). (f) Mutations in the interaction interface between MreC and RodZ disrupt their interaction as indicated by BTH assay. Pairs of plasmids expressing the indicated fusion proteins were co-transformed into BTH101. Spot test was performed as in (c). (g-h) Disruption of the interaction between RodZ and MreC results in cell shape defects. (g) Overnight cultures of Δ*rodZ* cells expressing the indicated form of RodZ^1-155^ were grown in LB, diluted in the same medium supplemented with 0.4% arabinose, and grown at 37°C for 2-3h. (h) Overnight cultures of strains SD464 expressing pSD315 and pSD315-AL-DD (P_206_::*mreC^AL-DD^-mreD)* were grown in LB, diluted in fresh medium supplemented with 60 µM IPTG after removal of arabinose, and grown at 37°C until OD_600_ reached about 0.4-0.6. Cells were collected, fixed and then imaged. Scale bar, 5 µm.

To test if RodZ^138-155^ mediated the interaction between RodZ and MreC, we employed the BTH assay. As expected, RodZ^1-155^ retained the ability to interact with MreC and MreB, while RodZ^1-138^ interacted with MreB but not MreC (Fig. 5c). Next, we conducted Co-IP experiments using cells expressing Flag-tagged MreC and GFP-tagged RodZ^1-155^ or RodZ^1-138^. A protein band matching the size of GFP-RodZ^1-155^ was detected in immunocomplexes isolated with an anti-Flag antibody in cells expressing both GFP-RodZ^1-155^ and Flag-MreC (Fig. 5d). However, GFP-RodZ^1-138^ was not detected in the immunocomplexes isolated with Flag-tagged MreC. In reciprocal tests, Flag-MreC was dectected in immunocomplexes isolated with anti-GFP antibody in cells expressing GFP-RodZ^1-155^ but not GFP-RodZ^1-138^ (Supplementary Fig. 9c). These results indicated that RodZ^138-155^ was critical for the interaction with MreC. In agreement with this, Alphafold 3 prediction of the RodZ-MreCD complex revealed that RodZ^138-155^ contacted the α-helical domain of MreC, which is kincked rather than a continuous helix shown in the resolved MreC structures (Fig. 5e). Mutating the putative contacting residues in RodZ^1-155^(Q140K,E141K) or MreC (A79D, L80D) abolished the interaction but not their interaction with other elongasome components in BTH test (Fig. 5f), suggesting that the mutated residues mediate the interaction between MreC and RodZ.

To test the importance of the RodZ-MreC interaction for elongasome function, we checked if disrupting it by the above mutations affected cell growth and shape. As shown in Fig. 5g and Supplementary Fig. 10a, RodZ^1-155^(Q140K,E141K) failed to support growth or restore rod shape of *ΔrodZ* cells when expressed from an arabinose-inducible promoter. Similarly, MreC(A79D,L80D)-MreD failed to complement and restore rod shape to MreCD depleted cells (Fig. 5h and Supplementary Fig. 10b). Western blotting showed that these mutations did not affect the stability of GFP-RodZ^1-155^ or GFP-MreC (Supplementary Fig. 10c-d). Also, BTH assay showed that MreC (A79D, L80D) did not affect MreC self-interaction (Fig. 5f). Thus, RodZ^138-155^ mediates RodZ’s interaction with MreC and is important for elongasome activity. We designated this domain Mre<u>C A</u>ctivating <u>D</u>omain(CAD).

### Multiple regions of RodZ are necessary for it to regulate elongasome activity

The discovery of the interaction between RodZ and MreC prompted us to further investigate the role of RodZ in regulating elongasome activity. Although RodZ is critical for maintenance of rod shape in wild type cells, the defect of *ΔrodZ* cells is largely suppressed by activating mutations in MreB, PBP2 and RodA (42), as well as in MreC as shown above. This argues that RodZ stimulates the PG synthetic activity of the elongasome. Previous domain analyses of RodZ showed that its cytoplasmic helix-turn-helix (HTH) domain is important for interaction with MreB and deletion of this domain reduced the ability of RodZ to maintain rod shape (25, 26). In line with this, a truncated form of GFP-RodZ lacking the HTH domain (RodZΔHTH) could not complement the Δ*rodZ* strain at low level of induction but did at high level of induction (Fig. 5b and Supplementary Fig. 9a). In addition to the HTH domain, a cytoplasmic and basic juxta-membrane (JM) domain of RodZ has been suggested to be essential for it to maintain rod shape, since replacing it with the cytoplasmic domain of MalF (MalF1-13), a typical integral membrane protein, abolished RodZ function (25). Interestingly, we found that the JM domain was not strictly required for its function, because replacing it with a sequence of similar length (RodZΔJM^2xMalF13^), but not shorter length (RodZΔJM^MalF13^), allowed RodZ to complement the *ΔrodZ* strain at higher levels of induction (Fig. 5b). Western blotting showed that neither replacement affected the protein level of GFP-RodZ (Supplementary Fig. 9b). Thus, the lack of function of (RodZΔJM^MalF13^) observed previously may be due to the substituted sequence being too short. These analyses indicates that the HTH, JM and CAD domains all contribute to RodZ function: at lower level of overproduction, RodZ variants lacking any one of the three domains display reduced ability to support cell growth and maintain rod shape, however, at higher level of expression, they can largely restore growth and rod shape to the Δ*rodZ* strain. These results complement a previous work from the de Boer lab, which showed that at least two of the three domains (HTH, JM and periplasmic (CAD) are necessary for RodZ to support rod shape formation(25).

### RodZ initiates cytoplasmic and periplasmic signaling cascades to activate elongasome activity

The above analyses suggest that the role of RodZ in elongasome function may be analogous to that of FtsN in regulating sPG synthesis, where FtsN activates sPG synthesis by interacting with FtsA and FtsQLBWI to switch them to the active state on both sides of the membrane (Fig. 1). It is possible that the HTH and JM domain of RodZ interact with the MreB cytoskeleton in the cytoplasm (52) and the CAD domain interacts with MreC in the periplasm, such that they are switched to the activate state to stimulate the synthetic activity of RodA-PBP2 (Fig. 6a). In this regard, the cytoplasmic region (HTH and JM domain) of RodZ could be termed the Mre<u>B</u> <u>A</u>ctivating <u>D</u>omain (BAD).

**Fig. 6.**
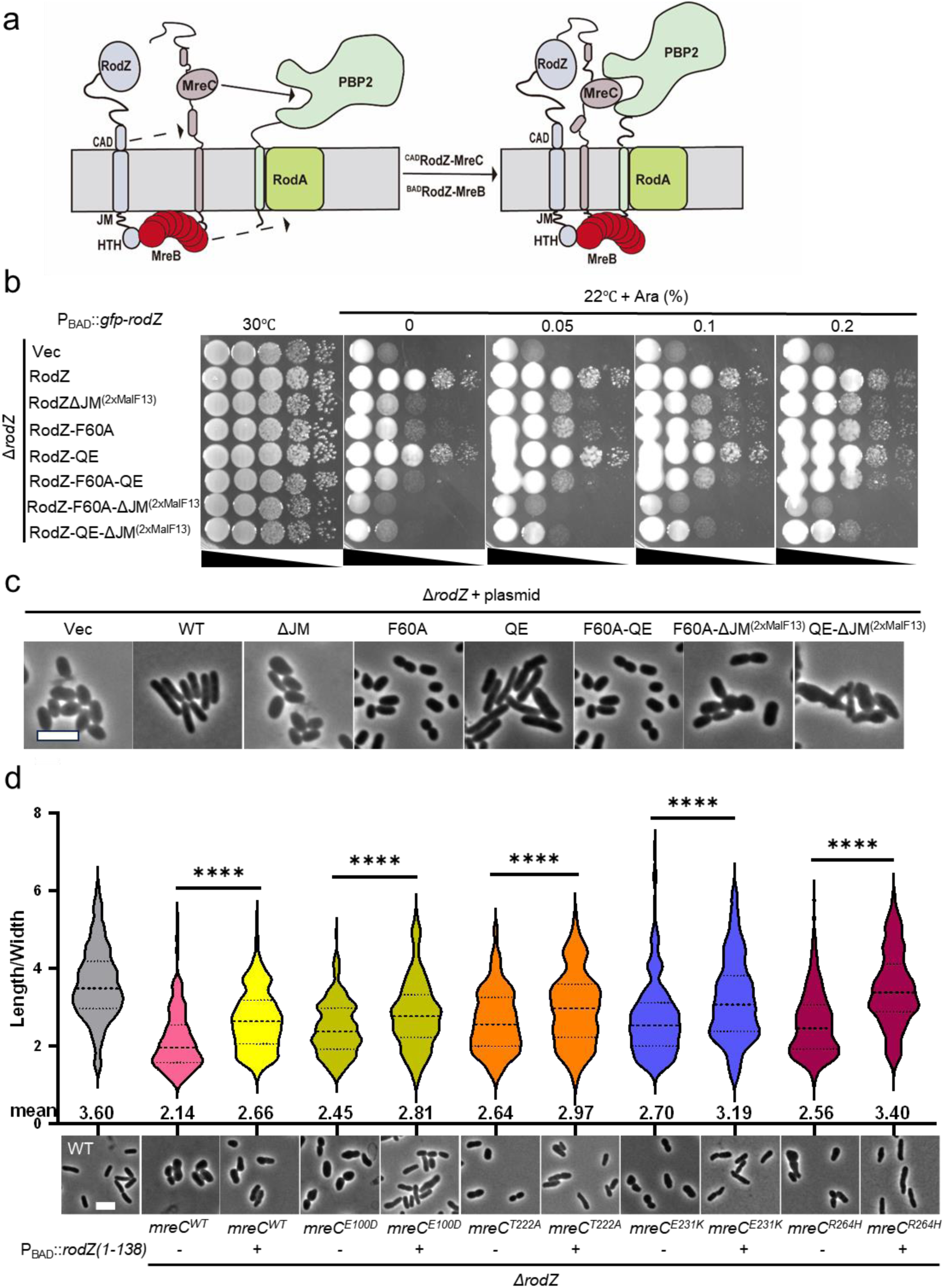
RodZ initiates cytoplasmic and periplasmic signaling cascades to activate elongasome activity. (a) A hypothetical model for the role of RodZ in regulation of elongasome activity. The periplasmic region of RodZ (138-155, CAD domain) interacts with MreC, inducing a conformational change in MreC so that it can bind to PBP2. Meanwhile, the HTH domain and JM domain (BAD domain) of RodZ binds to MreB, modulating its assembly and likely MreB’s interaction with the RodA-PBP2 complex. These two signaling cascades work together to activate the enzymatic activity of RodA-PBP2. (b-c) Both of RodZ’s interactions with MreB and MreC are important for cell shape maintenance. Overnight cultures of strain SD541 carrying plasmids pBAD18, pZ102(P_BAD_::*gfp-rodZ*), pZR106(P_BAD_::*gfp-rodZΔJM^(2xMalF13)^*), pZR102-F60A, pZR102-QE-KK, pZR102-QE-F60A, pZR106-F60A, pZR106-QE-KK, pZR103-F60A and pZR103-QE were grown in LB, diluted 1:100 into the same medium supplemented with 0.2% arabinose, and grown at 30°C until OD_600_ reached about 0.4-0.6. Then the cultures were diluted 1:20 in the same medium supplemented with 0.05% arabinose again, and grown at 30°C until OD_600_ =0.4-0.6. Cells were collected, fixed and then imaged. Scale bar, 5 µm. (d) Synergism of activating MreC mutations and RodZ^1-138^ in restoration of rod shape in *ΔrodZ* cells. Overnight cultures of strains carrying indicated *mreC* alleles (*mreC^E100D^*, *mreC^T222A^*, *mreC^E321K^* and *mreC^R264H^*) contained plasmid pBAD18 (empty vector) or pZR97 (pBAD18, P_BAD_::*rodZ^1-138^*) were diluted 1:100 in fresh LB medium with antibiotics and 0.4% arabinose (Ara), incubated at 30°C. Upon OD_600_ reaching 0.4 to 0.6, cells were fixed and imaged. The ratio calculated for each cell is plotted as a violin dot, showing the 25th and 75th percentiles, with the mean at the center, whiskers extend from -s.d. to +s.d. of the mean. Statistical significance was determined using an unpaired t test with Welch’s correction. Scale bar, 5 µm. ****P < 0.0001, two-tailed Student’s t-test. Number of cells analyzed (n≥200). Scale bar, 5µm.

To test the above model, we systematically analyzed GFP-RodZ mutants deficient in RodZ-MreB and/or RodZ-MreC interaction by expressing them from an arabinose-inducible promoter in a plasmid. As shown in Fig. 6b-c, introduction of the F60A mutation or replacement of the JM domain with 2×MalF13 to reduce the RodZ-MreB interaction decreased the ability of GFP-RodZ to restore cell growth and rod shape of *ΔrodZ* cells at lower level of arabinose concentration. However, a combination of them completely eliminated the ability of GFP-RodZ (GFP-RodZ^F60A^ΔJM^2xMalF13^) to support cell growth of the *ΔrodZ* cells and restore rod-shape, suggesting that the interaction between RodZ and MreB is critical for cell growth and shape maintenance. On the other hand, introduction of the QE/KK mutation in RodZ to disrupt the RodZ-MreC interaction did not noticeably affect the ability of GFP-RodZ to rescue growth and rod shape of *ΔrodZ* cells (Fig. 6b-c). Nevertheless, introduction of the mutation in the GFP-RodZ^F60A^ or GFP-RodZΔJM^2xMalF13^ background further diminished the ability of the mutants to complement, suggesting that disruption of RodZ-MreC interaction compromises RodZ function. Western blot of these GFP-RodZ mutants showed that they were expressed at similar level (Supplementary Fig. 11a), suggesting that the defects were not due to protein instability. It is notable that the QE/KK mutation strongly reduced the ability of GFP-RodZ^1-155^ to complement (Fig. 5g and Supplementary Fig. 9a), likely because the expression level of GFP-RodZ^1-155^ was lower than that of GFP-RodZ. Together, these results suggest that both RodZ’s interaction with MreB and MreC play important roles in cell growth and rod shape maintenance.

To further test the above model, we examined if RodZ^1-138^, which lacks the entire periplasmic region of RodZ, has a synergistic effect with activating MreC mutations in promoting rod shape formation. RodZ^1-138^ retains its interaction with MreB in the cytoplasm, thereby it should be able to initiate the activation signaling cascade in cytoplasm. However, as it cannot interact with MreC, it cannot trigger the signaling cascade in the periplasm. Activating MreC mutations partially restored rod-shape to Δ*rodZ* cells, its combination with RodZ^1-138^ may then reconstitute the cytoplasmic and periplasmic activation signaling cascades necessary for elongasome activity. To test this, we expressed RodZ^1-138^ from an arabinose-inducible promoter in a plasmid in Δ*rodZ* cells harboring activating MreC mutations. Note that expression of RodZ^1-138^ from the plasmid used in this experiment was unable to restore cell growth and rod shape of *ΔrodZ* cells (Fig. 6d and Supplementary Fig. 11b). As expected, expression of RodZ^1-138^ further improved the rod-shape morphology of Δ*rodZ* cells harboring activating MreC mutations (Fig. 6d), indicating that it enhanced elongasome activity independently of the MreC mutation. Taken together, these results indicate that the activating MreC mutations compensate for the loss of the RodZ-MreC interaction and that both the RodZ-MreB and RodZ-MreC interactions contribute to the optimal activity of the elongasome.

## Discussion

How the elongasome synthesizes lateral PG in rod-shaped bacteria remains incompletely understood. In this study, we employed genetic, biochemical and cytological approaches to investigate the mechanism governing elongasome activity. Our results reveal the roles of MreC(D) and RodZ in regulating elongasome activity and suggest a model in which RodZ initiates two signaling cascades, one in the cytoplasm through MreB and another in the periplasm via MreC(D), to stimulate the PG synthetic activity of RodA-PBP2. This model has similarities to that employed by FtsN to activate the divisome for sPG synthesis, implying an analogous regulatory mechanism for the elongasome and divisome.

Early studies of MreC suggested that it acted as an extracellular scaffold to organize lateral PG synthesis (53), but accumulating evidence indicates that it (in complex with MreD) is a regulator of elongasome activity (9, 34, 43, 44). First, it interacts with PBP2 directly and alters its conformation (44). Second, mutations in PBP2 and RodA that enhance elongasome activity largely suppress the growth and shape defects caused by dominant-negative (also inactive) MreC mutations in *E. coli* (9, 34), suggesting that the defects caused by inactive MreC variants are due to the lack of activity of RodA-PBP2. However, when RodA-PBP2 is constitutively active, the need of MreC could be largely bypassed (9). Structural analysis of MreC showed that it forms dimers or oligomers, which would mask its binding site for PBP2 (45–47). Interestingly, a recent study identified the γ domain of MreC as an important regulatory domain for its function as mutations within this region block elongasome activity, however, removal of the whole domain was tolerated (34). This observation suggests that MreC alternates between an active and an inactive state, with its γ domain playing a critical role in this transition. Putting together, these findings suggest that disruption of MreC self-interaction exposes the PBP2 binding site on MreC (switches it from the inactive state to the active state) such that it can interact with PBP2 to activate the RodA-PBP2 complex. However, whether this model is correct remains unknown.

In this study, we isolated a set of MreC mutations that confer resistance to the MreB antagonist A22. Many of these MreC mutations also partially suppress the defects of *ΔrodZ* cells (9). Moreover, these mutations serve as intragenic suppressors of a dominant-negative MreC mutation, R292H, which is believed to lock MreC in the inactive state (34). Thus, these MreC mutations share features with the activating mutations in PBP2 or RodA that result in a constitutively active elongasome (9), suggesting that they promote MreC in the active form. Along with previous findings on dominant-negative MreC mutations, these results lends support to the notion that MreC alternates between an inactive state and an active state. Some of the mutations reported in this study alter residues located in the MreC homo-dimer interface and disrupt MreC self-interaction or conformation, as indicated by results from the BMOE-mediated *in vivo* crosslinking assay. However, disruption of MreC self-interaction or conformation seems not a prerequisite for it to transition to the active state because the activating mutation R264H does not affect its self-interaction. Additionally, we found that the G156D mutation, as well as the R292H mutation which is located in the γ domain, reduced MreC self-interaction rather than increasing its self-interaction in BMOE-mediated *in vivo* crosslinking assay. Thus, it seems that there is no clear correlation between disruption of MreC self-interaction and its activity.

Although the effects of the activating mutations on MreC self-interaction vary, all tested ones appear to enhance the interaction between MreC and PBP2. In contrast, the inactive MreC mutations (G156D and R292H) reduce the interaction between MreC and PBP2. Furthermore, the regulatory γ domain of MreC appears to be necessary for the optimal interaction between MreC and PBP2. The absence of the γ domain completely eliminates the interaction between MreC and PBP2 in the BTH assay, and co-IP experiments showed that its absence partially reduced the interaction. Interestingly, introduction of the activating mutations into a truncated MreC variant without the γ domain (MreC^1-278^), restored the MreC-PBP2 interaction and alleviated the growth and shape defects caused by this mutant. This observation suggests that the γ domain is necessary for MreC to switch to the active state and its absence likely favors MreC in the inactive form.

MreD forms a tight complex with MreC and it likely acts together with MreC to regulate the activity of RodA-PBP2. Accordingly, we found that the combination of a MreC mutation and a MreD mutation can increase elongasome activity (confer A22 resistance), whereas either mutation alone could not, implying a coordinated action of the two proteins. A recent study showed that MreC and MreD from *Thermus thermophilus* form a dimer of heterodimer, in which the periplasmic membrane-proximal α domain of *^Tt^*MreC fits into a putative pocket of *^Tt^*MreD and interacts with additional periplasmic loops of *^Tt^*MreD (54). It is proposed that in the absence of MreD, the helical α domain of MreC exists in an extended conformation such that its β domain is too far above the membrane to interact with PBP2. However, upon binding to MreD, the α domain is shortened so that the β domain gets closer to the membrane to interact with PBP2 and activate the RodA-PBP2 complex (54). While this model is tempting, it should be noted that formation of the MreCD complex is likely not sufficient for MreC to interact with PBP2 effectively as MreC still needs to transition to the active state to interact with PBP2. As we shown in this study and discuss below, the most likely candidate to trigger MreC to interact with PBP2 is RodZ.

RodZ is conditionally essential for cell growth and maintenance of rod shape (25, 26, 42). The defects caused by *rodZ* deletion is readily suppressed by spontaneous mutations in MreB, RodA and PBP2 (42), rendering its role in the elongasome elusive. Interestingly, the *ΔrodZ* suppressor mutations, at least for the ones in RodA and PBP2, turn out to be activating mutations of the elongasome (9), indicating that the absence of RodZ results in reduced elongasome activity. Consistent with this, we showed that activating MreC mutations also partially suppress the growth and shape defect caused by *rodZ* deletion. Thus, RodZ may be an activator of the elongasome. In line with this, we identified a periplasmic region of RodZ (aa138-155, the CAD domain) that interacts with MreC directly and is important for elongasome function. Additionally, the cytoplasmic HTH domain and JM domain of RodZ and the activating MreC mutations promote rod shape formation synergistically. Taken together, our results suggest that RodZ stimulates elongasome activity through two parallel pathways. One pathway involves the regulation of MreB polymers via its HTH domain and JM domain, which we renamed as the BAD (Mre<u>B</u> <u>A</u>ctivating <u>D</u>omain) domain, while the other pathway involves a direct interaction between the CAD domain and MreC, which likely switches MreC to the active state (Fig. 7a). It is possible that in the absence of RodZ, neither MreB nor MreC is able to switch to the active form effectively, thus resulting in reduced elongasome activity and a partial loss of rod shape and conditional growth (Fig. 7b). However, in the presence of activating mutations in other elongasome components, such as MreB, MreC, RodA or PBP2, its role can be largely bypassed (Fig. 7c). We hypothesize that either one of the RodZ-induced signaling cascades can maintain rod shape if it is strong enough, for example, by overexpression of the RodZ truncations or activating MreB mutations. However, at the chromosomal level, both signaling cascades are important to fully activate the elongasome. It should be pointed out that the exact mechanism through which RodZ’s interaction with MreB leads to RodA-PBP2 activation remains unknown, but a previous study showed that a minimal elongasome consisting of only MreB and constitutively active RodA-PBP2 complex can compensate for the simultaneous loss of MreCD and RodZ, and partially maintains rod shape (9). Thus, there must be interactions between MreB and the RodA-PBP2 complex that await to be uncovered.

**Fig. 7.**
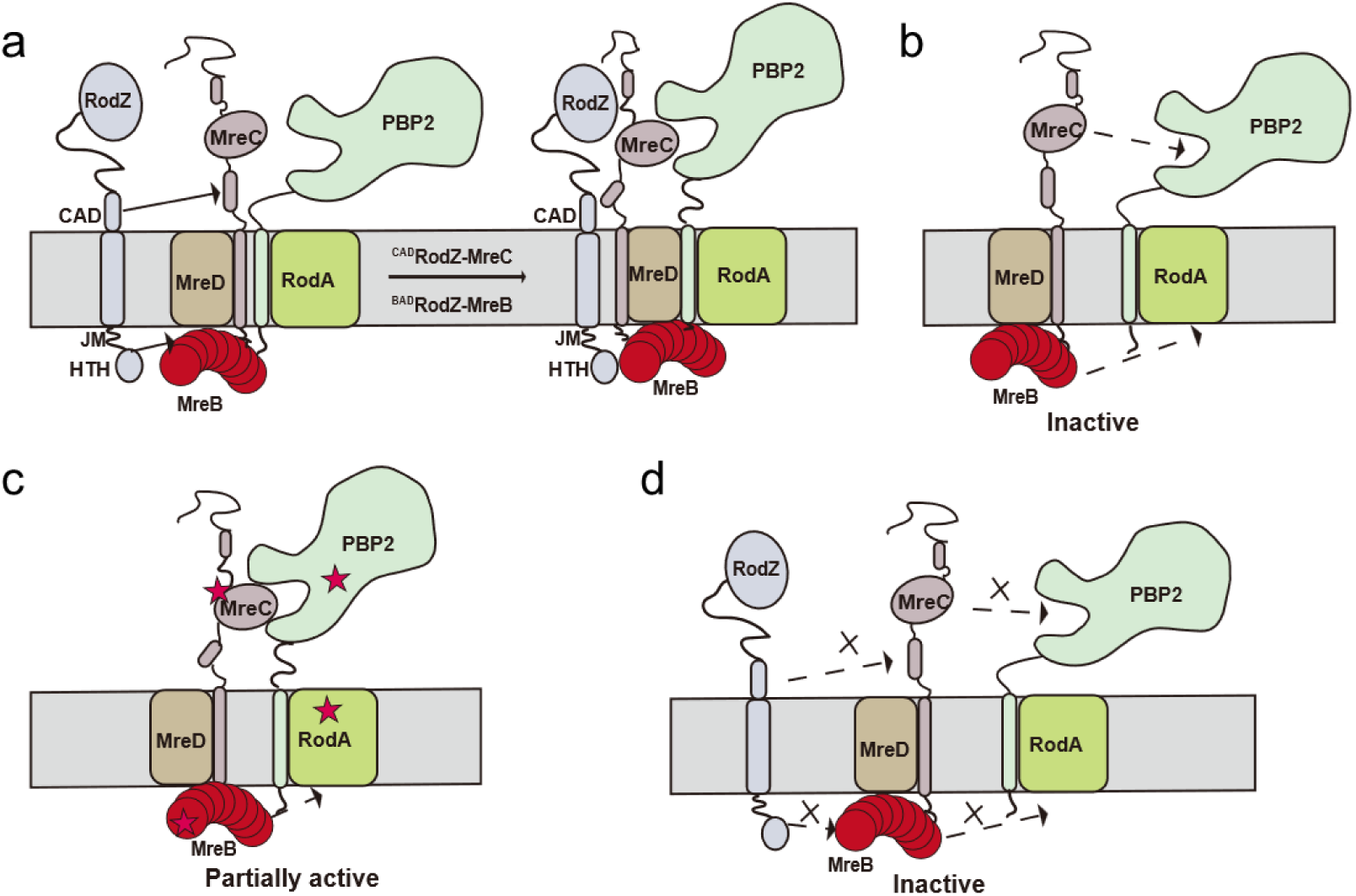
A working model for the regulation of elongasome activity. The elongasome is composed of six proteins: MreB, MreC, MreD, RodA, PBP2 and RodZ. MreB is the cytoskeleton in the cytoplasm, organizing the assembly of the complex and guiding the synthesis of lateral PG. RodA-PBP2 is the peptidoglycan synthase. MreCD functions as a direct regulator of the activity of RodA-PBP2, whereas RodZ functions as a trigger for activation of the elongasome. In wild type cells (a), the periplasmic Mre<u>C</u>-<u>A</u>ctivating <u>d</u>omain (CAD) of RodZ introduces a conformational change in MreC so that it can switch from the inactive state to the active state and interact with PBP2. The MreC-PBP2 interaction stabilizes the RodA-PBP2 complex in the active configuration, leading to the activation of the polymerase activity of RodA. Meanwhile, the HTH domain and JM domain (Mre<u>B</u>-<u>A</u>ctivating <u>D</u>omain, BAD) of RodZ modulates MreB assembly and initiates another signaling cascade in the cytoplasm, which can also activate the activity of RodA-PBP2. (b) In the absence of RodZ, neither MreB nor MreC is able to switch to the active state effectively, such that the activity of RodA-PBP2 is substantially reduced, resulting in a partial loss of rod shape and conditional growth. (c) Activating MreC mutations enhance the interaction between MreC and PBP2, leading to activation of RodA-PBP2, thereby partially compensating for the loss of RodZ. Activating MreB mutations can also partially compensate for the loss of RodZ, but how they work is unclear currently. Activating RodA and PBP2 mutations result in constitutively active RodA-PBP2 complex, bypassing the need of RodZ. (d) Disruption of either the cytoplasm or periplasm signaling cascade reduces the elongasome activity and results in cell shape defect.

Our findings reveal a similarity between the activation mechanism of the elongasome and that of the divisome. This similarity arises from numerous compositional and functional parallels between the proteins of the divisome and the components of the elongasome. Both complexes contain a related SEDS-bPBP pair (RodA-PBP2 *vs* FtsW-FtsI) that need to be activated, however, the regulatory components are unrelated (MreCD-RodZ *vs* FtsQLBN). Nonetheless, RodZ’s interaction with MreB and MreC in the cytoplasm and periplasm, respectively, is analogous to FtsN’s interaction with FtsA in the cytoplasm and FtsQLB in the periplasm. These interactions switch the regulators (MreB and MreCD for the elongasome, FtsA and FtsQLB for the divisome) to the active form. Consequently, these functional counterparts stabilize the SEDS-bPBP complexes in the active state to start PG synthesis.

## Methods

### Media, bacterial strains, plasmids and growth conditions

Cells were grown in LB medium (1% tryptone, 0.5% yeast extract, 0.5% NaCl and 0.05 g/L thymine) at indicated temperatures. Antibiotics were used at the following concentrations when necessary: ampicillin = 100 μg/mL; spectinomycin = 25 μg/mL; kanamycin = 25 μg/mL; tetracycline = 12.5 μg/mL; and chloramphenicol = 15 μg/mL. Strains, plasmids and primers used in this study are listed in Tables S2-4, respectively. Detailed procedures for construction of strains and plasmids are described in Supplemental Information with the primers listed in Table S4. Mutant alleles in the chromosome were introduced by allelic exchange and were moved between strains by phage P1-mediated transduction, and the antibiotic cassette was removed using FLP recombinase expressed from pCP20 (55).

### Mutagenesis

Mutagenesis was carried out by site-directed mutagenesis using the Quickchange II kit (Agilent) following the protocol provided with the kit or by overlap PCR. The primer pairs used for mutagenesis are provided in Table S4. All mutations were confirmed by sequencing.

### BTH assay

To detect the interaction between elongasome proteins or its mutants, appropriate plasmid pairs were co-transformed into BTH101. The next day, single colonies were resuspended in 1 mL LB medium, and 2.5 μL of each aliquot was spotted on LB plates containing 100 μg/mL ampicillin, 25 µg/mL kanamycin, 40 μg/mL X-gal and 100 μM IPTG. Plates were incubated at 30°C overnight before imaging.

### Allelic replacement in *E. coli*

Different alleles of *mreC* were introduced into the chromosome at its native locus by allelic replacement as described by the methods of Hamilton *et al* (50). The approach employs homologous recombination between a gene on the chromosome and homologous sequences carried on a plasmid temperature sensitive for DNA replication. The mutations were first introduced into a plasmid (pSC101^ts^, *mreC*) by site-directed mutagenesis or overlap PCR. These plasmids were transformed into the wild-type strain TB28 and selected on LB plates containing spectinomycin at 44°C.

Because the plasmids replicate at 30°C but not at 44°C, they integrated into the chromosome at the *mreC* locus, resulting in integrants carrying the mutations. Subsequently, these co-integrants were grown at 30°C to trigger a second recombination event, causing the excision of the plasmids. Depending on where the second recombination event took place, the chromosome would either have undergone a gene replacement or retain the original copy of *mreC*. 10-20 colonies were randomly picked for each desired *mreC* mutation and the *mreC* gene from these colonies was sequenced to detect whether the mutation was successfully introduced into the chromosome.

### Western blot

To measure the level of elongasome proteins and their mutants, overnight cultures of TB28 (wild type strain) harboring the respective plasmids were diluted 1:100 in LB medium with ampicillin and 100 μM IPTG. After growth at 37°C for 2 hours, samples were taken and adjusted according to the OD_600_ of cultures to make sure equal amount of samples was used for each strain. Cells were collected, resuspended in SDS-PAGE sample buffer and boiled for 10 min before they were loaded on the SDS-PAGE gel for analysis. Anti-GFP (TRANSGEN, HT801) and Anti-Flag (Proteintech Group, 66008-4-lg) antibody were used at a dilution of 1/10,000. Secondary antibodies were used at a dilution of 1/10,000.

### Immunoprecipitation

Overnight cultures of TB28 carrying the indicated expression plasmids were diluted 1:100 in 50 mL fresh LB medium, and cultivated at 37°C until OD_600_ reached 0.4-0.6. Cells were collected by centrifugation at 12,000 rpm for 10 min and resuspended in 1 mL PBST (1× PBS+0.5% Tween-20) containing 1% anti-protease cocktail (MCE) and lysed by sonication. The lysates were centrifuged at 12,000 rpm for 5 min to remove cell debris. 400 μL of the supernatant was added to pre-prepared Ab-coated magnetic beads and incubated overnight at 4°C. The bead-antibody-supernatant complexes were then separated using a magnetic rack and washed four times with 400 μL of PBST. Finally, the immunecomplexes were eluted with SDS-PAGE sample buffer and boiled for 10 min before they were loaded on the SDS-PAGE gel for western blotting analysis.

### Creation of MreC and MreD mutant library

*mreCD* was subjected to random PCR mutagenesis using the primer 5-EcoRI-mreC and 3-mreD-HindIII and plasmid pSD315 (pDSW210, P_206_::*mreC-mreD*). Gel purified PCR product was digested with EcoRI and HindIII and ligated into pDSW210 digested with the same enzymes. Ligation product was transformed into JS238 competent cells and plated on LB plates with ampicillin and glucose. About 30,000 transformants were pooled together and plasmids were isolated and stocked.

### Screen for A22 resistant MreC/MreD mutants

The control plasmid pSD315 or the mutagenized pSD315 (pSD315-M) was transformed into strain JS238. Transformants were then plated on LB plates containing IPTG (100 μM) and with or without A22 (10 μg/mL). Transformants of the control plasmid pSD315 could not grow on the plates containing A22 at 37°C because A22 was toxic to cells. Transformants of pSD315-M that could grow on the selective plates thus likely contained plasmids expressing MreC or MreD mutants that could suppress the growth defect caused by A22. Plasmids were isolated from A22-resistant colonies and transformed back into the parental strain JS238 to confirm that resistance was plasmid linked. The *mreC-mreD* operon from such plasmids were then sequenced to identify the mutation(s) conferring A22 resistance. In cases where the plasmid contained multiple mutations, the causative mutation was identified using site-directed mutagenesis or overlap PCR to generate single substitutions, followed by assessment of resistance to A22 by a spot test.

### BMOE-mediated *in vivo* cysteine cross-linking assay

A single or a pair of cysteine residues was introduced into a cysteine-free form of MreC based on the available dimer structure of *E. coli* MreC. Overnight cultures SD464 harboring pZR212 (pDSW210, P_206_::*flag-mreC^C175S,C199S^/mreD*) expressing the indicated alleles of *mreC* were diluted 1:100 in 5 mL LB with antibiotic and arabinose, and grew at 37°C for 2-3h, and then cultures were collected by centrifugation and washed twice with fresh LB to remove arabinose, resuspended in the same volume of LB supplemented with ampicillin, kanamycin, chloramphenicol and 60 µM IPTG, and grew at 37°C until OD_600_ reached 0.4 to 0.6. Cultures were taken and adjusted according to the OD_600_ of cultures to ensure equal amount of samples was used for each strain. Cultures were cooled down by placement on ice and spun at 10,000×g for 2 min at 4°C. The pellet was washed in ice-cold 2 mL PBSG (PBS, supplemented with 0.1% glycerol), spun at 10,000×g for 2 min at 4°C. The cell pellet was resuspended in 2 mL PBSG, supplemented with 5 mM EDTA. Half of the resuspended cells were subjected to the thiol-reactive crosslinker bismaleimidoethane (BMOE) at a concentration of 200 µM from a 20 mM stock in DMF, whereas the same amount of DMF was added to the other half of the cells as a control. After 15 min on ice, the reaction was quenched with 28 mM β-mercaptoethanol and spun for 10 min at 10,000 ×g at 4°C (Eppendorf table top centrifuge). The pellets were resuspended in SDS-PAGE sample buffer and boiled for 10 min before they were loaded onto the SDS-PAGE gel for analysis. Anti-Flag antibody (Proteintech Group, 66008-4-lg) was used at a dilution of 1/10,000.

### Microscopy: Phase contrast, epifluorescence

Phase contrast and epifluorescence images were collected on an Olympus BX53 upright microscope with a Retiga R1 camera from QImaging, a CoolLED pE-4000 light source and a U-Plan × Apochromat phase contrast objective lens (100×, 1.45 numerical aperture [NA], oil immersion) using VisiView software. Fluorescence images of GFP were acquired using the Chroma EGFP filter EGFP/49002. For ^SW^MreB-mNeon, the exposure time for wild-type strain was about 500 ms.

### Microscopy: Observations of ^SW^MreB-mNeonGreen localization in *E. coli*

To investigate the impact of A22 treatment on the localization of ^SW^MreB-mNeonGreen in MreC mutants, strains (RZ15 to RZ21) expressing *^SW^mreB-mNeonGreen* (from attλHC897) were grown to log phase. Subsequently the cultures were diluted 1:100 in 5 mL fresh LB medium and IPTG was added to a final concentration of 200 μM and grown at 30°C for about 2 h and 2 µg/mL A22 was added to the culture (designated as time 0). Cells were harvested at 0 min, 40 min and 120 min after the addition of A22 and immobilized on 2% LB-agarose pads for imaging.

### Microscopy: Observations of cell morphology in *E. coli*

#### MreCD depletion strain

Overnight cultures of strain SD464 (TB28, *mreCD<>aph*/pSD352 (P_BAD_::*mreCD*)) carrying plasmid pDSW210 (P_206_::*gfp*) or pSD315 (P_206_::*mreCD*) and its derivatives carrying different alleles of *mreC* were diluted1:100 in fresh LB medium with antibiotics and 0.2% arabinose, grown at 37°C for 2 h. Cells were then collected by centrifugation and washed twice with fresh LB and resuspended in the same volume of LB medium. These arabinose-free cultures were diluted 1:100 in fresh LB medium with antibiotics, grown at 37°C for 2-3 h. Then cultures were diluted 1:10 in fresh 5 mL LB medium again and IPTG was added to a final concentration of 60 μM and grown at 37°C about 3-4 hours, cells were immobilized on 2% agarose pad for photograph.

### *ΔrodZ* strain

To test the impact of activating *mreC* mutations on Δ*rodZ* cells, overnight culture of the *ΔrodZ* strains containing the indicated *mreC* alleles at the native genomic locus (E100D, P150L, S153R, D154N, P182L, Q184L, T222A, E231K and R264H) were diluted1:100 in fresh LB medium with antibiotics and grown at 30°C for 3 h. Then the cultures were diluted 1:10 in fresh LB medium again until OD_600_ reached 0.4-0.6. Cells were concentrated by centrifugation and immobilized on 2% LB-agarose pads before imaging.

To verify the impact of *rodZ* mutations on Δ*rodZ* cells, overnight culture of strain SD541 (TB28, *yhdE<>cat* Δ*rodZ<>aph*) carrying plasmid pZR102 (pBAD18, P_BAD_:*:gfp-rodZ*) and its derivatives were cultured as above and induced with 0.4% arabinose. 3 hours after induced by arabinose, samples were harvested and immobilized on 2% LB-agarose pads for imaging.

To test whether activating MreC mutants collaborate with RodZ^1-138^ to restore rod shape in Δ*rodZ* cells, plasmids pBAD18 (empty vector) or pZR97 (pBAD18, P_BAD_::*rodZ^1-138^*) were transformed into strains harboring indicated *mreC* alleles at the native genomic locus (SD541, SD542, SD548, SD549, and SD550), respectively. The above strains were cultured as above and induced with 0.4% arabinose. Cultures were grown at 30°C for 3-4 hours until the OD_600_ reached 0.4-0.6. Cells were concentrated by centrifugation and immobilized on 2% LB-agarose pads before imaging.

To assess the impact of impaired MreC-RodZ and MreB-RodZ interactions on the restoration of rod shape in *ΔrodZ* cells, overnight cultures of strain SD541 (TB28, *yhdE<>cat* Δ*rodZ<>aph*) harboring plasmids expressing *gfp-rodZ* and its variants were diluted 1:100 into fresh medium supplemented with 30 µM IPTG and incubated at 30°C until the OD_600_ reached approximately 0.4-0.6. Subsequently, the cultures were diluted 1:10 again into the same medium containing 30 µM IPTG and further incubated at 30°C about 2h. Cells were concentrated by centrifugation and immobilized on 2% LB-agarose pads before imaging.

### Quantification and statistical analysis

Cell lengths and widths were measured by ImageJ. Briefly, phase contrast images were imported into ImageJ, converted to 8-bit versions with scale bar embedded. Each cell was outlined and measured. Cells exhibiting partial view were excluded. The aspect ratio was calculated by dividing the length measurements by the width measurements. The data were plotted in GraphPad Prism (v.10.0.1) and statistical analysis of aspect ratio done in GraphPad Prism using a parametric unpaired t test assuming gaussian distribution but not equal SD (Welch’s correction). Images were cropped in ImageJ. More than 200 cells were analyzed for each strain.

## Data availability

All strains and plasmids used in this study are listed in Table S2-3, respectively. All data presented in this study will be publicly available as of the data of publication. Source data are provided with this paper.

## Acknowledgments and funding sources

We thank members of the Du lab, Chen lab and Lutkenhaus lab for advice and helpful discussions to carry out this study This study was supported by National Natural Science Foundation of China (grant 32070032 and 32270049 http://www.nsfc.gov.cn/), the Fundamental Research Funds for the Central Universities (grant 2042021kf0198), and the Young Top-notch Talent Cultivation Program of China to S.D.

## Competing interest statement

The authors declare no competing interests.

